# Structurally divergent and recurrently mutated regions of primate genomes

**DOI:** 10.1101/2023.03.07.531415

**Authors:** Yafei Mao, William T. Harvey, David Porubsky, Katherine M. Munson, Kendra Hoekzema, Alexandra P. Lewis, Peter A. Audano, Allison Rozanski, Xiangyu Yang, Shilong Zhang, David S. Gordon, Xiaoxi Wei, Glennis A. Logsdon, Marina Haukness, Philip C. Dishuck, Hyeonsoo Jeong, Ricardo del Rosario, Vanessa L. Bauer, Will T. Fattor, Gregory K. Wilkerson, Qing Lu, Benedict Paten, Guoping Feng, Sara L. Sawyer, Wesley C. Warren, Lucia Carbone, Evan E. Eichler

## Abstract

To better understand the pattern of primate genome structural variation, we sequenced and assembled using multiple long-read sequencing technologies the genomes of eight nonhuman primate species, including New World monkeys (owl monkey and marmoset), Old World monkey (macaque), Asian apes (orangutan and gibbon), and African ape lineages (gorilla, bonobo, and chimpanzee). Compared to the human genome, we identified 1,338,997 lineage-specific fixed structural variants (SVs) disrupting 1,561 protein-coding genes and 136,932 regulatory elements, including the most complete set of human-specific fixed differences. Across 50 million years of primate evolution, we estimate that 819.47 Mbp or ~27% of the genome has been affected by SVs based on analysis of these primate lineages. We identify 1,607 structurally divergent regions (SDRs) wherein recurrent structural variation contributes to creating SV hotspots where genes are recurrently lost (*CARDs*, *ABCD7*, *OLAH*) and new lineage-specific genes are generated (e.g., *CKAP2*, *NEK5*) and have become targets of rapid chromosomal diversification and positive selection (e.g., *RGPDs*). High-fidelity long-read sequencing has made these dynamic regions of the genome accessible for sequence-level analyses within and between primate species for the first time.

## INTRODUCTION

An early and still unmet grand challenge of the Human Genome Project has been to reconstruct the evolutionary history of every base pair of the human reference sequence^1–5^. To do so requires both a diverse sampling of nonhuman primate (NHP) genomes but also a more complete assembly of those genomes so that all forms of variation can be assessed without bias introduced from a superior quality reference^6–13^. Early attempts to sequence closely related ape species focused primarily on characterizing simpler forms of variation (e.g., single-nucleotide variants, (SNVs)) from portions of the genome that could be readily aligned to human^7–10,13^. As long-read sequence assemblies began to emerge, our ability to catalog larger forms of structural variation significantly improved resulting in a series of more contiguous NHP genomes. These new references, however, represented “squashed” assemblies where allelic variation was collapsed and the most complex forms of gene-rich structural variants (SVs) were still not resolved, including recently duplicated sequence^14–19^. Advances in long-read sequencing technology over the last three years now allow for most of these regions to be accurately sequenced and assembled to a degree where both paralogous and allelic variation can be readily distinguished^20–23^. Numerous studies focused on the human lineage have shown that such regions are incubators for the emergence of new genes, adaptive evolution while also contributing to disease, and disease susceptibility^24–26^.

To better characterize SVs and these complex genic SV regions, we generated genome assemblies of eight NHP genomes using two long-read sequencing platforms. Our plan was twofold: First, we wanted to broaden the phylogenetic diversity by sequencing additional NHP genomes using the same sequencing platform (in this case continuous long-read sequencing or PacBio CLR) that had been initially applied to the other ape references to minimize sequencing technology biases. This included sequence and assembly of primate genomes representing gibbon (*Nomascus leucogenys*), marmoset (*Callithrix jacchus*), and one owl monkey (*Aotus nancymaae*) (Table 1). Second, we wanted to leverage the higher accuracy and assembly contiguity of HiFi (high-fidelity) sequencing data by sequence and assembly of all NHP genomes where haplotypic differences could be distinguished. These served as a means to validate all fixed structural variation events as well as provide complete haplotype-resolved access to any particular regions of interest without the need to construct and annotate these different NHP genomes for yet a third time.

**Table 1.** Primate genome sequence and assembly.

| Common name | Scientific name | Individual ID | Sex | CLR raw data and assembly |  |  | HiFi raw data and assembly |  |  | Iso-Seq (Gbp) | ONT (Gbp) |
| --- | --- | --- | --- | --- | --- | --- | --- | --- | --- | --- | --- |
|  |  |  |  | reads (coverage) | assembly (contig N50, Mbp) | QV | reads (coverage) | assembly (contig N50, Mbp) | QV hap1/hap2 |  |  |
| Chimpanzee | *Pan troglodytes* (common chimpanzee) | Clint_PTR | M | 117 | 12.27 | 39.19 | 37 | 66.89 /49.98* | 45/44 | 1.94 | 294 (178*) |
| Bonobo | *Pan paniscus* (pygmy chimpanzee) | Mhudiblu_PP A | F | 74 | 15.06 | 39.25 | 39 | 50.45 /36.22* | 47/47 | 1.38 | 124* |
| Gorilla | *Gorilla gorilla gorilla* (western lowland gorilla) | Kamilah_GGO | F | 84.3 | 9.52 | 38.72 | 31 | 38.19 /37.87* | 46/46 | 1.84 | 264* |
| Orangutan | *Pongo abelii* (Sumatran orangutan) | Susie_PAB | F | 94.9 | 11.07 | 34.83 | 43 | 62.38/ 58.39* | 42/42 | 1.09 | 272 (126*) |
| Gibbon | *Nomascus leucogenys* (northern white-cheeked gibbon) | Asia_NLE | F | 92.5* | 12.78* | 38.65 | 31* | 44.67 /34.99* | 43/43 | 15.25 * | 97* |
| Macaque | *Macaca mulatta* (Rhesus monkey) | AG07107_M MU | F | 66 | 46.61 | 36.18 | 29 | 18.81 /19.01* | 51/52 | 104.5 8 | 329 (231*) |
| Marmoset | *Callithrix jacchus* (white-tufted-ear marmoset) | CJ1700_CJA | F | 66* | 25.23* | 42.95 * | 39* | 103.97 /87.06* | 58/58 | 18.43 * | NA |
| Owl monkey | *Aotus nancymaae* | 86718_ANA | F | 56.3* | 9.85* | 37.4* | 31* | 55.92 /44.99* | 57/57 | NA | 91* |

\* New data in this study

## RESULTS

### Genome assembly of NHP genomes

Building on our previous analysis of African great ape genomes^14,17,19^, we first sequenced and assembled three additional female NHP genomes using CLR sequencing, namely, white-cheeked gibbon (*Nomascus leucogenys*), the common marmoset (*Callithrix jacobus*), and owl monkey (*Aotus nancymaae*) Each genome was sequenced to high depth (>56-fold coverage), assembled, and error corrected as described previously^14,16,17,19^ (Supplementary Figure 1 and Supplementary Table 1). We generated highly contiguous (contig N50=9.9 to 25 Mbp) squashed assemblies of ~2.84-2.9 Gbp with an overall sequence accuracy of >99.98% (Table 1 and Supplementary Table 1). Next, to further reduce sequencing error and increase our ability to investigate more complex regions, we sequenced the same eight NHP samples using PacBio HiFi sequencing^17,27^ (Table 1; Supplementary Figure 2 and Supplementary Table 1). We used hifiasm to produce haplotype-resolved genomes that were substantially smaller among monkeys (5.84 to 6.23 Gbp, diploid) when compared to nonhuman apes^21^ (6.12 to 6.98 Gbp). These HiFi assemblies are estimated to be more accurate (QV=42 to 58 or 99.9937% to 99.9998% accuracy) and significantly more contiguous (contig N50=19 to 104 Mbp) when compared to the CLR draft genome assemblies (Table 1 and Supplementary Figure 3).

### NHP sequence divergence and incomplete lineage sorting (ILS)

As a baseline for sequence divergence among the lineages, we mapped the HiFi sequence data from each NHP back to human and computed single-nucleotide divergence (Methods). The mean autosomal sequence divergence ranged from 1.3% to 9.83%, consistent with the expected phylogeny, and was predictably higher than that of the X chromosome (0.99% to 8.24%; Figure 1a and 1b, Supplementary Table 2). We note that these estimates are also slightly higher than earlier reports likely because a great fraction of repetitive DNA is being included among NHPs^8,19^. For example, among the apes ~92% of the human genome is aligned in contrast to the New World monkey lineages where 64% and 59.7% of the sequence from owl monkey and marmoset are unambiguously aligned (Supplementary Table 3). An assembly-based comparison yields similar results but involves a smaller fraction of the genome due to extensive and more complex forms of structural variation (Supplementary Figure 4 and Supplementary Table 3).

**Figure 1.**
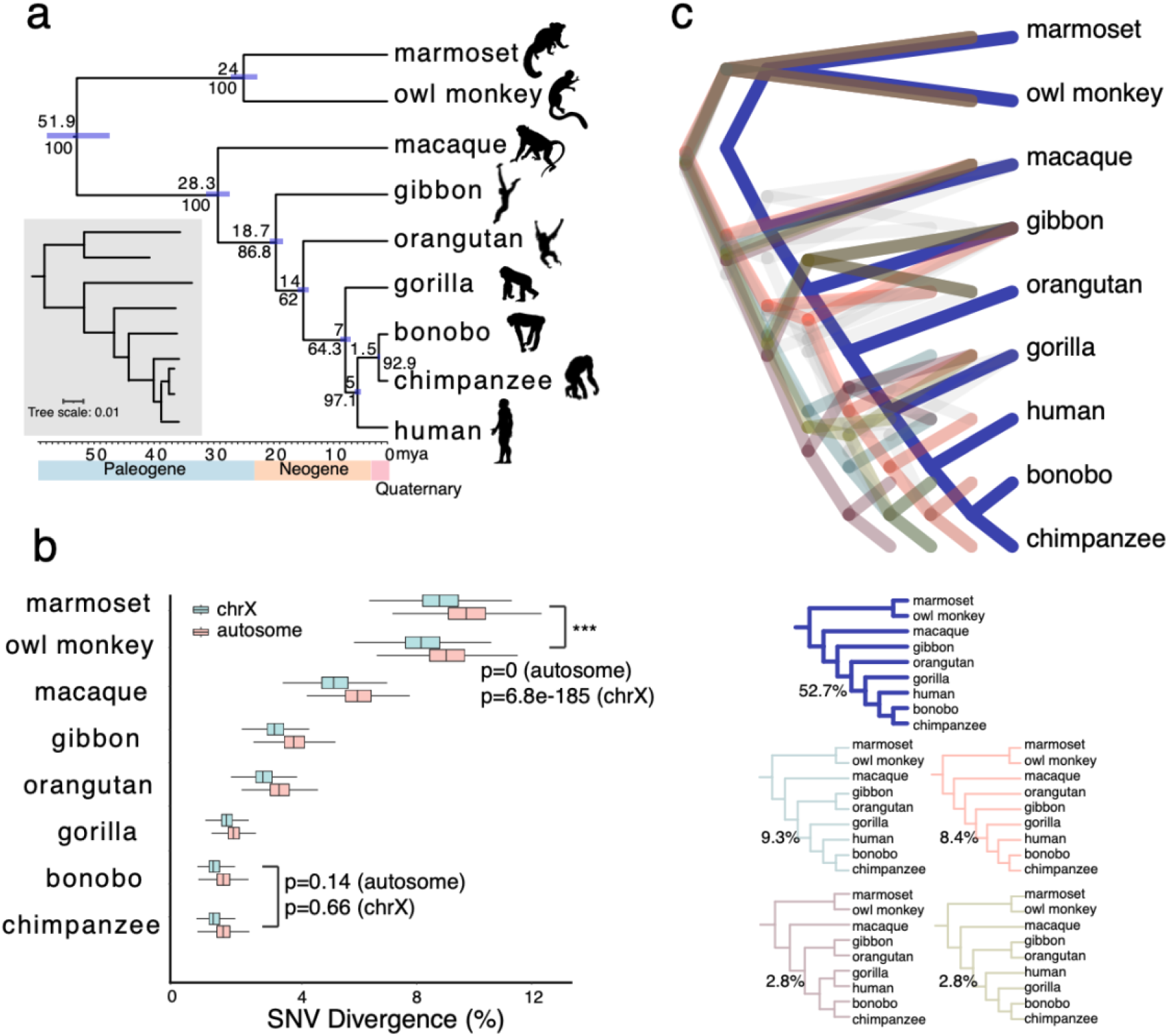
Primate phylogeny and SNV divergence between NHPs and humans. **(a)** A primate time-calibrated phylogeny was constructed from a multiple sequence alignment (MSA) of 81.63 Mbp of autosomal sequence from nine genomes. The estimated species divergence time (above node) with 95% confidence interval (CI, horizontal blue bar) was calculated using BEAST2. All nodes have 100% posterior possibility support, and the gene tree concordance factor (gCF) is indicated (below node). The inset (gray) depicts a maximum likelihood phylogram generated using IQ-TREE2, which reveals a significantly shorter branch length in owl monkey, with respect to marmoset. **(b)** SNV divergence calculated by mapping HiFi sequence reads to human GRC38 separately for autosomes and the X chromosome (excluding pseudoautosomal regions). Approximately 85% of the genome was aligned for Old World monkey and apes and ~60% for New World monkey. The owl monkey shows significantly less divergence compared to human than the marmoset (Wilcoxon rank sum test). An analysis using 20 kbp nonoverlapping segments from the assembly gives almost identical results (Supplementary Figure 4). **(c)** The percent of trees showing an alternate tree topology are indicated (percentages are drawn from a total of 302,575 gene trees): 159,546 (52.7%) support the primate topology depicted in panel a.

We used these data to generate a time-calibrated phylogeny for the nine primate species, including human (Figure 1a and 1b; Supplementary Tables 4-6). We constructed more than one million complete multiple sequence alignments (MSAs) at a resolution of 500 bp (518.9 Mbp of aligned sequence). While the majority of trees (52.7%) are consistent with the generally accepted phylogeny, the fraction of alternate topologies is, once again, greater than previous estimates^9,13,17,28^ (Figure 1c, Supplementary Table 4). Most of the difference can be attributed to potential ILS during African ape or great ape speciation as gene tree concordance factors show the lowest values in these two nodes (gene tree concordance=64.3 and 62, respectively)^29^. Lineage-specific branch lengths are generally balanced with one notable exception: the owl monkey branch length is significantly shorter and divergence to human significantly lower when compared to marmoset (Figure 1a). An analysis of 16,244 gene trees using human as an outgroup to both owl monkey and marmoset shows that the owl monkey evolves significantly slower (p=0 autosome, p=6.85·10^-185^ for the X chromosome) (Supplementary Figure 5). Excluding potential sites of ILS, we estimated split times of the species and find that mean split times of the apes better match the lower bounds of previous estimates^30–36^ (Supplementary Table 7).

### Primate lineage-specific versus shared SVs

We applied a three-pronged approach to discover and validate SVs (≥50 bp) mapping to the euchromatic portion of the primate lineages^37,38^. Using read-based and assembly-based callers (pbsv, Sniffles and PAV), we first compared the eight NHP genomes against the human reference genome, including three additional human genomes (CHM13, HG00733 and NA19240) to mitigate the effect of human polymorphism and missing variants in a particular reference (Supplementary Table 8). In total, we identified 2.23 million putative insertions and 1.89 million deletions in these nine lineages. Using both HiFi sequence data and genome assemblies, we validated 1.85 million insertions and 1.63 million deletions (mean validation rate: 86.79% and 89.37%, respectively) (Supplementary Table 9). We note that genome-based HiFi and CLR SV calling are highly congruent (>95%) although HiFi tended to recover larger insertions (Supplementary Figure 6). Finally, we generated Oxford Nanopore Technologies (ONT) data from the same primate DNA samples and manually inspected a subset (900 SV events) for confirmation using this orthogonal sequencing platform estimating a false positive rate and a false negative rate of ~2.6% and 11.4%, respectively (Supplementary Table 10).

To distinguish fixed from polymorphic events, we further genotyped (Methods) the validated SVs against Illumina whole-genome sequence (WGS) data from a panel of 120 genomes (30 humans and 90 NHPs, Supplementary Table 11)^39–43^. We projected the 1,338,997 fixed events (441,453 deletions and 897,544 insertions) onto the primate phylogeny (Figure 2a; Supplementary Tables 12 and 13) classifying events as shared or lineage-specific^17^ (Methods). The number of SV events correlates strongly with evolutionary genetic distances separating species (Figure 2b) with characteristic insertion peaks at ~6 kbp and 300 bp—full-length *L1* and *Alu* mobile element insertions (Supplementary Figure 7 and Supplementary Table 14). Remarkably, we estimate that 27.2% of the genome (819.47 Mbp) has been subjected to structural variation across these nine lineages with fixed insertions outnumbering deletions approximately two to one (the total length of shared and lineage-specific insertions is ~524.8 Mbp versus ~294.68 Mbp of deletions) (Figure 2a). The excess of insertions is greatest for the ancestral ape and African great ape lineages (~2-to 3-fold) (Figure 2a and Supplementary Table 13) and this twofold excess is still observed when calibrating for the number of fixed SNV differences^44,45^ (Figure 2b; Supplementary Figures 8 and 9).

**Figure 2.**
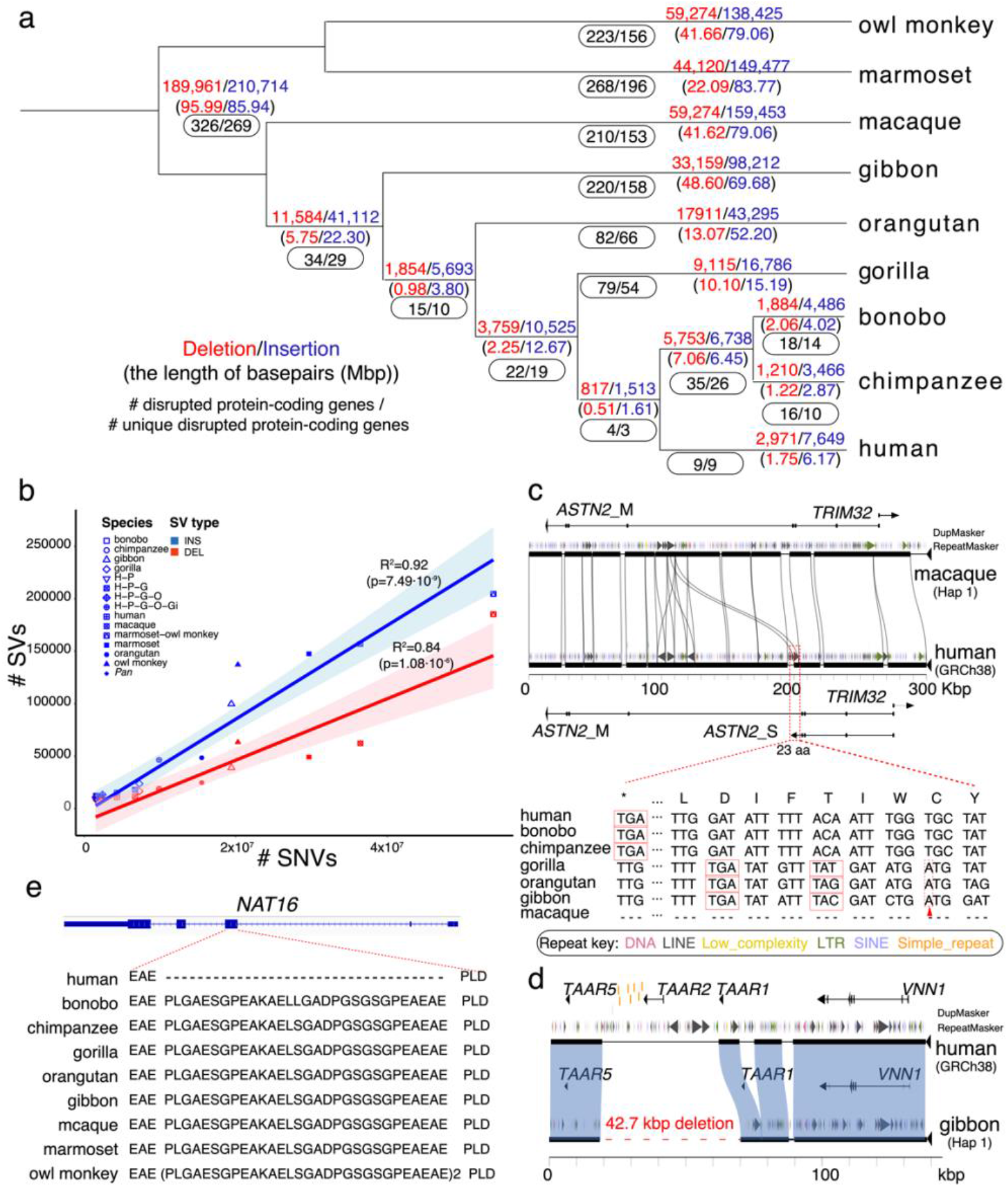
Primate genome structural variation. **(a)** The number of fixed structural variants (SVs) including deletions (red) and insertions (blue) are shown for each branch of the primate tree (number of events above the line and number of Mbp below). The number of “disrupted” protein-coding genes based on human RefSeq models are also indicated (black oval) with the total number of events (first number) and the subset specific to each lineage (second number). **(b)** The number of fixed SVs correlates with the accumulation of SNVs in each lineage (comparison to GRCh38) for both deletions (red) and insertions (blue). **(c)** An ape-specific fixed *L1* insertion (shown with a red dashed line box) in the human genome but not in the macaque genome (Miropeats alignment) serves as an exapted exon of the short isoform of astrotactin 2, *ASTN2*, in human. The coding sequences of the exon are shown in the bottom panel. The red triangles represent 1 bp insertion resulting in a frameshift in gorilla, orangutan, and gibbon. The red box represents the stop codon. **(d)** A 42.7 kbp lineage-specific deletion in the gibbon genome (red dashed line) deletes *TAAR2* and seven enhancers (shown in orange) compared to the human (GRCh38) (Miropeats comparison). **(e)** A 90 bp deletion (30 amino acids) human-specific deletion of *NAT16* (NM_001369694) removes 30 amino acids in humans compared to all other NHPs.

A small fraction of fixed primate SVs affect genes (~18.78 Mbp of deletions and ~1.31 Mbp insertions). Using human gene annotation as a guide, we annotated the fixed SVs against the human gene models (GRCh38, RefSeq) and the regulatory element database (ENCODE V3) with Variant Effect Predictor (VEP)^46,47^. These fixed SVs intersect 6,067 genes, including 1,561 protein-coding genes, and 136,932 regulatory elements. The latter includes 2,389 promoter-like (PLS) and 16,455 proximal enhancer-like signatures (pELS) potentially disrupted by 16,671 fixed SVs (Supplementary Table 15). We estimate that 244 genes and 1,759 regulatory elements are novel and several are likely to confer functional effect (Supplementary Figures 10 and 11). Such is the case for the 3,741 bp *L1PA5* insertion shared in apes mapping to the last exon of the neuronal-function gene, astrotactin 2 (*ASTN2*), which encodes a glycoprotein that guides neuronal migration during the development of the central nervous system^48,49^. The insertion creates a novel transcript isoform resulting in a new exon in human (NM_1884735) and this innovation is accompanied by a 1 base-pair deletion in this exon, which in gibbon, orangutan, and gorilla is incapable of read-through due to a frameshift from the reciprocal 1 bp insertion (Figure 2c and Supplementary Figure 12). Similarly, the Aggrecan (*ACAN*) gene, important in stature and brachydactyly in humans^50^, has been altered in the great ape lineage by a 60 bp deletion, which eliminates part of the chondroitin sulfate attachment domain (Supplementary Figure 13). In gibbons, we identify a large ~42.7 kbp deletion of the neurogenesis-associated gene, trace-amine associated receptor 2, (*TAAR2*) along with seven of its enhancers (Figure 2d and Supplementary Figure 14). Loss of this brain-expressed gene in knockout mice has been shown to result in higher levels of dopamine and lower levels of norepinephrine in the striatum and hippocampus respectively^51^. A complete list of these gene and gene-regulatory fixed SVs is provided along with additional discussion (e.g., *AR*, *SPATA1, ELN*, and *MAGEB16*) (Supplementary Tables 16 and 17, Supplementary Figures 15-18, and Supplementary Discussion).

We also reassessed human-specific changes and the effect of potential reference biases in discovery. Importantly, 7,169 human-specific SVs have been reclassified, in part, because of the inclusion of more outgroup species in addition to the use of more accurate sequence aligner (minimap2 vs. blasr) that improves alignment within repetitive regions such as subtelomeres^52,53^ (Supplementary Figures 19 and 20). Nevertheless, we identified 13 additional genes and 252 additional regulatory elements as potentially disrupted compared to our previous report^19^ (Supplementary Figures 21 and 22). This includes, for example, a 90-base pair deletion within the third exon of N-acetyltransferase 16 (*NAT16*) resulting in 30 amino acid loss in human lineage with respect to all other NHPs. The event was confirmed in all humans by genotyping and by full-length transcript sequencing (Figure 2e and Supplementary Figure 23). *NAT16* is highly expressed in the brain and pituitary and is responsible for Nα-acetylhistidine synthesis, but its biological function remains unknown.

To assess the effect of using a human reference genome to classify such events, we repeated ape-specific SV analyses using an assembled African human genome and the orangutan ape, instead as the reference genomes to base the comparison. As expected, the analyses reclassified approximately 34 gene-disruption events and led to a reduction of SVs most notably with respect to insertions (Supplementary Figure 24). For example, using orangutan as a reference reduces the number of lineage-specific insertions in orangutan (56,389 vs. 77,933), chimpanzee (2,020 vs. 4,471), bonobo (3,108 vs. 5,886), and human (13,446 vs. 16,696) lineage-specific insertions (Supplementary Figures 25 and 26, Supplementary Table 18). The intersect of these two sets provides the most conservative set of lineage-specific changes on each branch. Consistent with the previous analyses, we find that the number of insertions is ~2-3 times than that of deletions.

### Structurally Divergent Regions (SDRs)

In addition to increased accuracy and haplotype resolution, another major advantage of HiFi-based assemblies is their 4-to 6-fold increase in sequence contiguity (Table 1). During our comparison of monkey and ape chromosomes, we identified much larger, structurally divergent regions (SDRs) that had been missed or incompletely assayed by our standard SV analyses (Supplementary Figures 27 and 28). These regions were often gene-rich but had eluded complete characterization due to their sequence divergence and/or structural complexity^54^. We, therefore, developed a graph-based approach to more systematically identify such regions (>10 kbp in length) in apes and macaques that could not be readily mapped to the complete human genome (T2T-CHM13) with >85% sequence identity^55^. We identified 1,704 SDRs and validated 1,607 SDRs using two independent approaches^56^ (validation rate: 94.3%; Methods) (Supplementary Tables 19 and 20). SDRs were large (average 127.4 kbp, Supplementary Figure 28) and enriched 3.6-fold for large segmental duplications (SDs) (Supplementary Figure 29; p=0). Specifically, 423 SDRs (26.3%) contain at least 10 kbp of annotated SDs while 1,184 appeared to map to relatively unique regions of the genome (Supplementary Tables 19 and 20), although subsequent sequence analysis identified 2.2% of these (1.07 Mbp) as lineage-specific SDs not present in human SD annotations.

Similar to the SVs, we genotyped all SDRs using Illumina WGS from primate population samples^39–43^ and successfully assigned 1,050 of the SDRs to lineage-specific branches on the primate phylogeny (Supplementary Figure 30). However, 557 SDRs show evidence of recurrent or serial SVs among multiple NHP lineages and the majority (62.3% or 347/557) associate with SD sequences (Supplementary Table 19). We constructed a null model for the distribution of SDRs and identified 184 distinct hotspot regions where we predict significant large-scale and recurrent structural variation among different primate lineages (331 recurrent SDRs) (Figure 3a and Supplementary Table 21). Of these, 88% (162/184) harbor SDs and 56% (103/184) of these hotspots correspond to 631 genes, including many known medically relevant regions such as *CFHR, RHD, LPA, APOL, AMY1* and the Major histocompatibility complex (MHC) locus (such as *MICA/MICB* and the complement *C4/C3* genes)^57–60^ (Figure 3a and Supplementary Figures 31-33). Others are completely novel or have been partially described based on analyses of specific primate genomes^57–60^. A gene ontology analysis predicts an expected enrichment for the known widespread loss of olfactory receptors in primates (p=3.10^-85^) but also genes associated with other biological processes, including thiol-dependent ubiquitinyl hydrolase activity (p=1.9·10^-24^), antimicrobial activity (p=2.2·10^-5^), innate immune response (p=5·10^-5^), neurotransmitter receptor activity (p=2.5·10^-4^), etc. (Supplementary Table 22). Notably, most of these enrichments are associated with core duplicons including *DEFBs*, *NPIPs*, *RGPDs*, *CYPs*, *NBPFs*, *GOLGAs*, *UGTs*, *RHDs*, and *USPs*^60,61^ (Supplementary Table 23).

**Figure 3.**
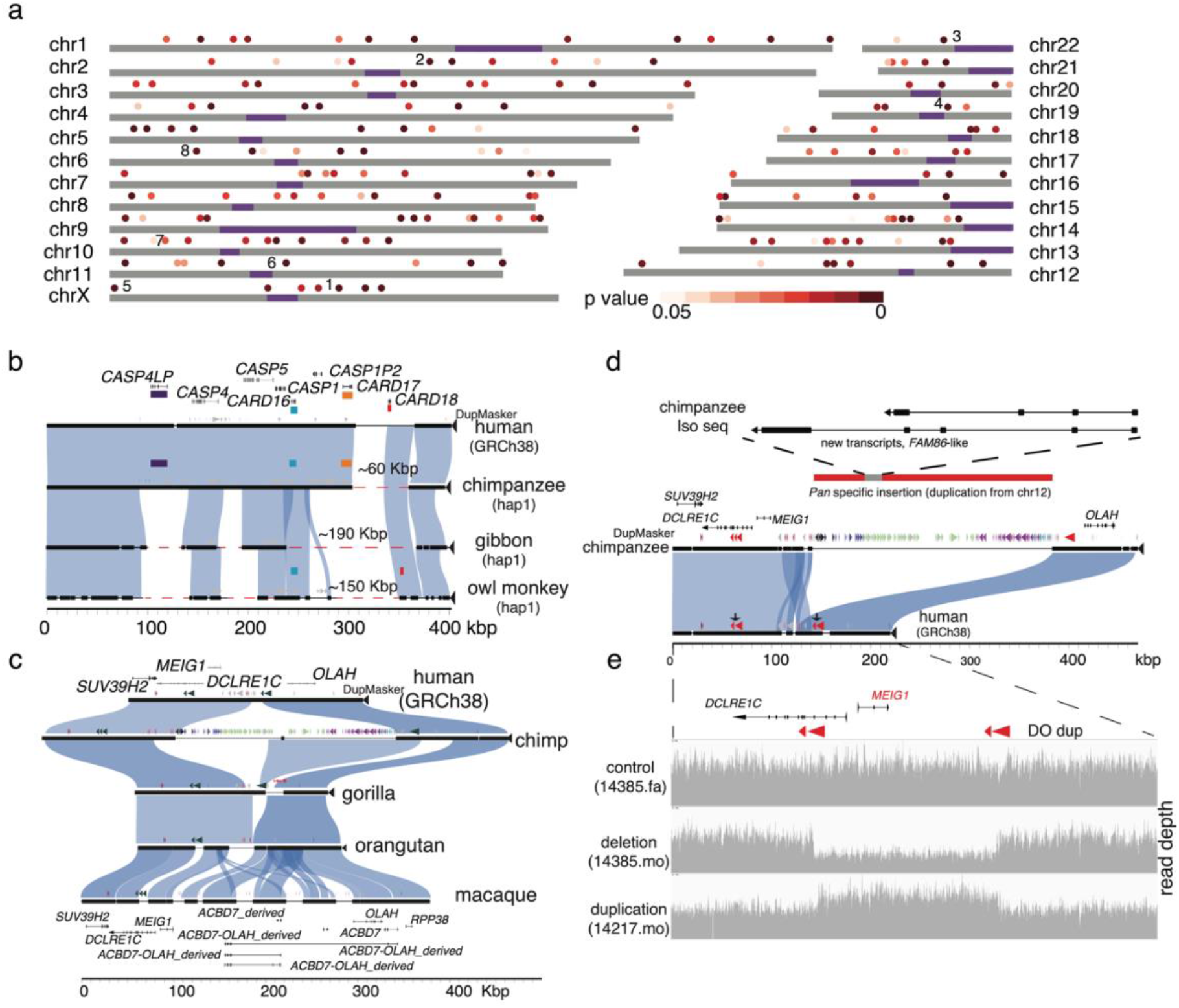
Structurally divergent regions (SDRs) of the primate genome. **(a)** A schematic of human chromosomes (T2T-CHM13) depicts SDR hotspots where recurrent rearrangements occur in excess. Heat map indicates significance based on simulation model (dark (p=0) to light red (p=0.05)). Centromeres are depicted in purple. Enumerated regions identify specific gene families or regions of biomedical interest (1: *UPRT*, 2: *RGPDs*, 3: *USP41*, 4: *ZNFs*, 5. *IL3RA_2*, 6: *CARDs*, 7: *OLAH*, and 8: *MHC*). **(b)** Recurrent deletion of the caspase recruitment domain (*CARD*) gene family. SafFire plot (https://github.com/mrvollger/SafFire) shows a ~58 kbp deletion of *CARD18* (orange) in the *Pan* lineage, multiple deletions (~190 kbp total) in gibbon of *CARD16* (blue), *CARD17* (red) and *CARD18*, and multiple deletions ~150 kbp, including *CARD17* (red), in marmoset. **(c)** SafFire plot of SDR mapping to genes *OLAH, MEIG1*, and *ABCD7* in human shows a large ~250 kbp insertion of segmental duplications (SDs; colored arrowheads) in chimpanzee within the intergenic region between *MEIG1* and *OLAH. OLAH* is deleted in gorilla by an independent lineage-specific deletion (~30 kbp). Multiple independent insertion events in macaque add ~190 kbp of sequence, including a duplication of *OLAH* in macaque. Full-length transcript sequencing of macaque using Iso-Seq supports the formation of five novel transcripts, including four *OLAH-ABCD* fusion events and a derived *ABCD7* (macaque gene models below). **(d)** The chimpanzee-specific 250 kbp SD from chromosome 12 creates a novel multi-exonic gene model supported by Iso-Seq transcript sequencing in chimpanzee (upper panel) with an unmethylated promoter (Supplementary Figure 36). The insertion simultaneously deletes one of two directly orientated (DO) SDs in chimpanzee. **(e)** In humans, the DO repeats associate with the breakpoints of recurrent deletions and duplications of the spermiogenesis gene *MEIG1*. Two females carrying a deletion and a duplication (as measured by sequence read depth) are depicted from a population sample of 19,584 genomes (CCDG, https://ccdg.rutgers.edu/). The carrier frequencies for microdeletion and microduplication in control samples are 0.026% and 0.189%, respectively.

A few examples of these hotspot regions are illustrative. We confirmed, for example, that the *CARD18* (caspase recruitment domain family member 18) was lost in the ancestral *Pan* lineage by ~60 kbp deletion event^7^. We identified, however, a larger and independent deletion of ~190 kbp in the gibbon lineage that completely removes the entire gene cluster— *CARD16* (pLI=0.04), *CARD17* (pLI=0), and *CARD18* (pLI=0.05). A third independent deletion of ~150 kbp removed yet another member, *CARD17*, in the owl monkey suggesting that this entire gene family has been under relaxed selection during primate evolution (Figure 3b and Supplementary Figure 34). Other hotspots are more complex, such as the *OLAH-ACBD7* region showing evidence of both gain and loss of genes (Figure 3c). In gorilla, *OLAH* (pLI=0) is deleted by a ~32 kbp deletion (Supplementary Figure 35) whereas in macaque the locus has been the target of ~190 kbp duplication that truncates *OLAH* in that lineage but also creates a new copy of *ACBD7*, which is actively transcribed as a fusion gene (Figure 3c). In *Pan*, the same region has been the target of a ~250 kbp SD that originated from chromosome 12 and produces a *Pan*-specific transcript with an open-reading frame (ORF) of 97 amino acids whose promoter region is hypomethylated (Figure 3d, Supplementary Figure 36). This large insertion of an SD in the *Pan* lineage also had the benefit of removing one of two directly orientated duplications flanking *MEIG* (meiosis/spermiogenesis associated 1), theoretically eliminating recurrent microdeletion/microduplication of *MEIG1* in the *Pan* lineage (Figure 3d and 3e). A 28 kbp genomic duplication region has been depleted in orangutans, but this has not resulted in any alteration of gene content (Supplementary Figure 37). *MEIG1* (pLI=0.05) is a spermiogenesis-related gene and *MEIG1* deficiency severely disrupts mouse spermatogenesis and is potentially associated in human infertility^62–64^.

In order to test the potential for SDRs to serve as cradles for gene innovation, we repeated our SDR analysis in a more distantly related primate. Using our graph-based approach, we compared human and marmoset and identified 697 SDRs (~38.45 Mbp) that could not be orthologously aligned to the complete human reference genome. Next, we manually clustered them into 270 distinct SDR events since these two genomes are too divergent (Supplementary Table 24). For the purpose of gene discovery, we also generated ~5.13 million full-length cDNA transcripts from 10 distinct primary tissues from the common marmoset (Table 1 and Supplementary Table 25). We identified five regions that showed evidence of novel or structurally divergent transcripts that lacked orthologous counterparts in the human genome (Supplementary Figure 38 and Supplementary Table 24). Of particular interest was a gene-rich region of human chromosome 13 that had been subject to a series of inversions and duplications increasing by ~350 kbp in size and adding nine putative marmoset-specific genes (Figure 4a). We searched for gene expression using the marmoset Iso-Seq transcript resource and confirmed expression for five of these—*VPS36, UTP14C, NEK5, THSD1*, and *CKAP2* broadly in the brain as well as other tissues (Figures 4b, Supplementary Figure 39 and Supplementary Table 26). In addition, our phylogenetic analysis estimates that the marmoset-specific duplication of the *THSD-NEK* region occurred ~11.9 million years ago (mya) and these duplicated genes maintain a protein-encoding ORF with numerous amino acid replacements as well as changes in gene structure when compared to progenitor copies (Supplementary Figure 40).

**Figure 4.**
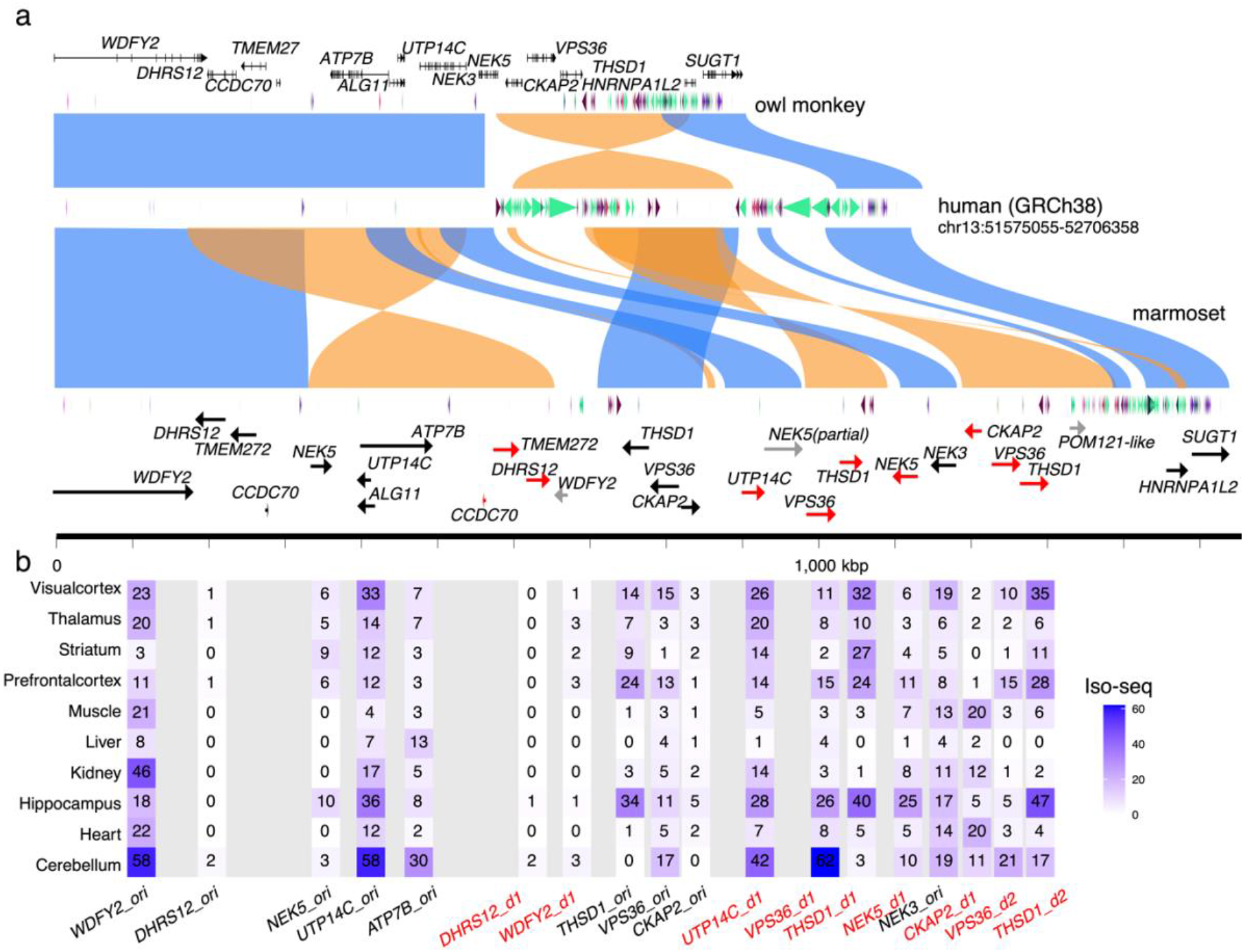
Marmoset-specific genes in a SDR. **(a)** SafFire plot comparing the organization of a gene-rich region of ~1.1 Mbp in human (middle), owl monkey (top), and marmoset (bottom) genomes. Human and marmoset differ mainly by a large 250 kbp inversion (orange) associated with the addition of 150 kbp of SD at the boundary of the inversion in humans (colored arrowheads). The corresponding region in marmoset has expanded by ~400 kbp due to inversion and marmoset-specific SDs creating marmoset-specific paralogs (red arrows) of *CCDC70, TMEM272, DHRS12, UTP14C, THSD1, VPS36, NEK5* and *CKAP2*. **(b)** Iso-Seq full-length non-chimeric transcript sequencing from 10 marmoset primary tissues confirms transcription of 8/10 of the paralogous copies and the maintenance of an open-reading frame in at least six of these marmoset-specific gene candidates.

### Recurrent RGPD duplications and restructuring of ape chromosome 2

Our SDR analysis of apes identified five SDRs on human chromosome 2 associated with a single core duplicon: *RGPD* (chr2:105859737-114023252, p=0) (Figure 5a). Core duplicons were previously described as actively transcribed gene families associated with the expansion of interspersed SDs in the human–ape lineage^61^. In particular, *RGPD* is a fusion gene/transcript formed by the duplication and juxtaposition of the two ancestral genes *RANBP2* and *GCC2* less than 15 mya^65^. Given the contiguity of the HiFi genome assemblies, we focused on a detailed reconstruction of the evolutionary history of this gene family across a ~7 Mbp region of chromosome 2 relating its expansion to large-scale structural changes and potential gene innovation associated with the SDRs in humans (Supplementary Figures 41 and 42). No evidence of *RGPD* genes exist in macaques, marmosets, or owl monkeys where only the ancestral *RANBP2* and *GCC2* genes are found syntenically among all primates (Supplementary Figure 42). Phylogenetic analyses confirm its formation and general expansion in copy number in the ancestral ape lineage (Supplementary Figure 43). Both the phylogeny and the sites of integration, however, reveal that most interspersed duplications are independent—the result of recurrent SDs or gene conversion events (Figure 5a and 5b, Supplementary Figure 43). For example, none of the gibbon or orangutan duplicate copies map syntenically to each other or other African great apes—thus, although orangutan has multiple *RGPD*s, all originated independently and none have orthologs among the other apes and group as distinct clade within the tree (Figure 5b and Supplementary Figure 43). We identify only one paralogous gene, *hRGPD2*, that is syntenic and orthologous among the African great apes. Within the five different ape lineages, we estimate ~20 independent mutation events (total length: ~1.2 Mbp) representing one of the most extreme examples of homoplasy (Figure 5a and Supplementary Figure 42).

**Figure 5.**
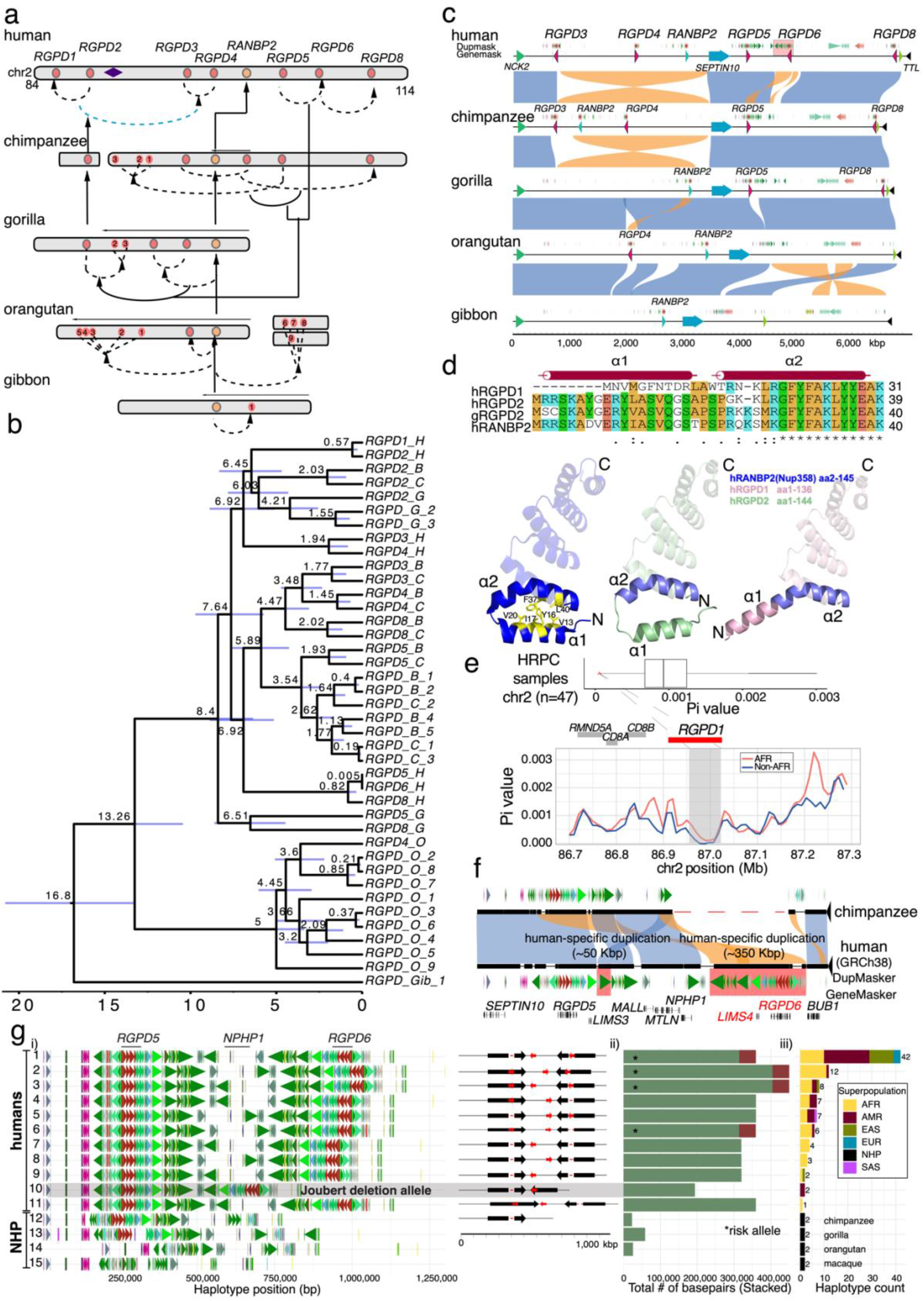
Evolution, selection, and disease susceptibility of the *RGPD* gene family. **(a)** Schematic depicting *RGPD* genes (red dots) compared to its progenitor gene *RANBP2* (orange dot) in human, chimpanzee, gorilla, orangutan, and gibbon. Shared ancestral copies among the lineages are indicated (vertical arrows) in contrast to lineage-specific duplications (black) or gene conversion events (blue dashed arced arrow). The majority of copies have expanded in a lineage-specific fashion in each ape lineage. **(b)** A maximum likelihood tree based on a 58.98 kbp MSA of 40 *RGPD* great ape copies outgrouped with a sole gibbon copy. Nodes are dated with BEAST2 with the mean age of divergence shown above the node (95% CI blue bar) for human (H), bonobo (B), chimpanzee (C), gorilla (G), orangutan (O), and gibbon (Gib) copies. The analysis confirms lineage-specific expansion with all nodes receiving 100% posterior possibility. **(c)** A comparison of ~7 Mbp on chromosome 2 among ape genomes showing that large breakpoints in synteny (colored rectangles) often correspond to sites of *RGPD* SD insertions (blue arrows). **(d)** Human genetic diversity (pi) calculated in 20 kbp windows (slide 10 kbp) from 94 haplotype-resolved human genomes (HPRC) for a 700 kbp region of chromosome 2. A segment mapping to the human-specific gene *RGPD1* shows the lowest genetic diversity on chromosome 2 (top panel, red arrow) in haplotypes of both African (red) and non-African (blue) descent. The data suggest that the *RGPD1* region may have been under recent selection in the ancestral human population. **(e)** AlphaFold predictions of the protein N-terminus structure RANBP2 (blue), h*RGPD1* (pink), and h*RGPD2* (green) predict that differences in amino acid composition alter the secondary structure of two alpha helices (α1 and α2) in the human-specific *RGPD1* copy. The X-ray crystal protein structure of hRANBP2 (Nup358, PDB: 4GA0) confirms that the α1 and α2 interface is maintained as a result of critical hydrophobic amino acids located in the N-terminus. Specific amino acid changes in hRGPD1 break the hydrophobic interface between α1 and α2 but not in the ancestral hRGPD2 or RANBP2 predicting the emergence of a human-specific protein structure. **(f)** SafFire plot (top panel) comparing the chimpanzee genome and human highlights the formation of a 350 kbp human-specific duplication creating *RGPD6* (red shading). **(g)** Analysis of 94 human haplotypes shows that the *RGPD6* locus is largely fixed among all humans but that the organization of the flanking SDs differs significantly. We identify 11 distinct structural haplotypes in the human population predicting both disease susceptibility as well as protective haplotypes for nonallelic homologous recombination (NAHR). NAHR between inverted repeats (large black arrows) predisposes to recurrent inversion of the region while NAHR between directly orientated repeats (red arrows) deletes the *NPHP1* allele creating the pathogenic allele associated with juvenile nephronophthisis and milder forms of Joubert syndrome^67^. This predisposition to disease, thus, arose as a result of the emergence of human-specific duplication of the *RGPD* gene family.

Most of the *RGPD* interspersed SDs were accompanied by both local restructuring of the duplication blocks as well as larger scale structural rearrangements of the chromosome 2 flanking sequence especially in association with large-scale inversions in different NHP lineages (Figure 5c and Supplementary Figure 44). Haplotype-resolved sequence assemblies allowed the origin and spread of lineage-specific copies to be distinguished phylogenetically (Figure 5b). Human *RGPD3* and *RGPD4* are not phylogenetically, for example, orthologs of chimpanzee *RGDP3* and *RGPD4* even though they appear syntenic (Figure 5b and Supplementary Figure 43) suggesting potential gene conversion. In addition, the emergence of many *RGPDs* in apes appears to have been driven by recurrent large-scale inversions, duplicative transpositions, and deletions within a ~7 Mbp genomic region over the last 15 million years of evolution creating unique configurations and distinct copies in each ape lineage (Supplementary Figure 44).

*RGPD1* is a human-specific paralog predicted to have arisen ~570 thousand years ago (kya) within the *Homo* lineage at ~0.57 mya (Figure 5b). This specific copy has several amino acid replacements at the protein N-terminus with respect to all other human RGPDs—this change is predicted to alter the protein structure between hRGPD1 and its antecedent hRGPD2^66^ (Figure 5d). In this regard, it is interesting that the *hRGPD1* genomic region shows a dearth of genetic diversity based on the analysis of Human Pangenome Reference Consortium (HPRC) haplotype-resolved assemblies (pi value=4.65•10^-5^, p<0.05, TajimaD= −1.98) (Figure 5e and Supplementary Figure 45) consistent with the region potentially being subjected to a selective sweep specifically and recently in the human lineage.

In comparison to human, most of the copies mapping to bonobo and chimpanzee chromosome 2 represent independent expansions from ancestral *RANBP2* that also gave rise to human *RGPD5, RGPD6*, and *RGPD8* (Supplementary Figure 43). Of note, *RGDP6* is a human-specific gene copy that arose via segmental duplication or gene conversion from human *RGPD5* most recently (~5.2 kya, 95% CI [0.002,16.08]) (Figure 5b). The interval between these human-specific copies, which includes *NPHP1*, is subjected to both inversion toggling and microdeletion associated with Joubert syndrome and juvenile nephronophthisis as a result of nonallelic homologous recombination (NAHR) between inverted and directly orientated duplications^67^’^69^, respectively (Figure 5f and Supplementary Figure 46). We examined 94 human phased haplotypes from the HPRC and Human Genome Structural Variation Consortium^38,69–71^ and identified 11 distinct structural configurations—four predisposing to microdeletion (Figure 5g; Supplementary Figures 46-50 and Supplementary Table 27). We also identified as single pathogenic allele deleting *NPHP1* (HG00733) and confirmed maternal transmission (Supplementary Figures 51-53). A maximum likelihood phylogenetic analysis identified the most closely related (non-deleted) haplotype and breakpoint analysis confirms that the deleted allele arose from one of the haplotypes predisposing to microdeletion (Supplementary Figure 51). Given the recent evolutionary restructuring of this region of chromosome 2, it follows that this predisposition to microdeletion is specific to the human lineage.

## DISCUSSION

Using three long-read sequencing platforms across multiple primate genera, we present a comprehensive analysis of SVs within euchromatic DNA of the primate order^15,19^. The use of HiFi data and inclusion of additional NHP species as well as genotyping in population samples significantly improves earlier surveys of fixed SV events^39–43^ and extends the analysis deeper within the primate phylogeny. Among the great apes for example, we identify 13 genes and 1,759 regulatory elements not previously reported^19^ (Supplementary Figures 21 and 22). The addition of other primate genomes identified lineage-specific SDR events in the gibbon (n=680), macaque (n=219), and marmoset (n=697) lineages (Supplementary Figure 30). Similarly, while we identify all 16 previously identified ape-specific genic SVs; 13/16 are no longer classified as (great) ape-specific SVs (Supplementary Table 28) due to the inclusion of other NHP lineages^15^. Finally, the use of a highly contiguous orangutan genome as an alternate reference, helped reduce earlier human genome reference biases by refining and polarizing the set of fixed SVs that occurred specifically since humans diverged from the other ape lineages (Supplementary Table 18). Among the 6,067 genes (both coding and noncoding) and 136,932 regulatory DNA associated with fixed SVs, we find a significant enrichment in transcription regulation (p=1.1·10^-9^), sensory transduction (p=6.3·10^-3^), cell division (p=2.3·10^-2^), and vocal learning (3.4·10^-3^) (Supplementary Table 29). These data serve as a rich resource for the characterization of gene expression differences and candidate mutations for adaptation among NHPs.

The overall topology of the primate phylogenetic tree is consistent with previous expectations with the proportion of ILS generally increasing as more of the repetitive content is accessed by long-read sequencing technology^17^ (Figure 1). Our comparison of two New World monkeys lineages, however, reveals significant acceleration of the marmoset SNV branch length when compared to that of the owl monkey (branch length: 0.024 vs. 0.017). This finding is also consistent with the shorter blocks of synteny in the marmoset lineage when compared to the human genome (only 102 regions >500 kbp compared to 169 regions >500 kbp in the owl monkey) and the significant increase in the number of recent SDs (165.7 Mbp in marmoset vs. 125.7 Mbp in owl monkey) (Supplementary Table 30). The slower evolution of the owl monkey lineage compared to marmoset may simply be a consequence of differences in reproductive longevity as has been proposed^40^ or changes in the generation time of the two lineages during evolution. The three major clades of New World monkeys, however, are thought to have diverged over a short time frame (19-24 mya)^35,36,72,73^ (Figure 1a). Studying multi-generational pedigrees, Thomas and colleagues showed a 32.5% reduction in the rate of *de novo* mutation in owl monkey when compared to that of apes with an overall mutation rate of 0.81·10^-8^ per site per generation^40^. Our results suggest that this reduced mutation rate may be longstanding property of the *Aotinae* with the net consequence that the owl monkey genome is less derived when compared to marmoset. These findings have some practical considerations regarding the use of these different New World monkeys as models for human disease^74–76^.

The greater accuracy afforded by HiFi sequencing allowed more complex regions of genetic variation to be assembled contiguously across the primates (e.g., MHC). We developed a graph-based approach to systematically identify 1,604 SDRs among apes and macaque (Figure 3) of which a third (n=557) showed evidence of recurrent structural variation and were enriched for SDs. We hypothesize that these hotspots of recurrent structural variation and their associated 631 genes (mean pLI=0.133) demarcate either regions of the ape genome no longer under selection (e.g., *CARD18, OLAH*, etc.) or regions where rapid structural diversification has facilitated the emergence of new genes showing signatures of positive selection (e.g., *RGPD, NPIP, NPF*)^77–79^ (Figure 5) and/or important for adaptive specializations in different primate lineages^24,80,81^. Ironically, the innovations often come at a cost with respect to fitness as the SDRs are associated with human disease susceptibility regions (e.g., 1q22.3, 2q13, 16p11.2, 10p13), such as the human-specific duplication of *RGPD5* and Joubert syndrome deletion alleles (Figure 5).

Our analysis also suggests that SDRs are common in the primate genome though with few exceptions these regions have not been considered as part of previous large-scale sequencing efforts because of 1) difficulties in their assembly and 2) challenges they pose in alignment even among closely related species when fully resolved. We identified, for example, SDRs in marmoset compared to owl monkey giving rise to marmoset-specific duplicate genes (Figure 4). Using our resource of ~5.13 million full-length transcripts, we show that these duplicate genes are expressed in the brain, maintain an ORF, and emerged specifically since marmoset diverged from other owl monkey ~20 mya (Supplementary Figure 54). The ancestral genes have critical functions: *NEK5*, for example, is member of NimA family of serine/threonine protein kinases involved in cell differentiation while *CKAP2* (cytoskeleton associated protein 2) is involved in cell division^82,83^. These findings caution against simply using human gene models to annotate NHP genomes or to assess NHP gene expression differences from single-cell RNA sequencing experiments. Understanding the gene innovations in such previously inaccessible complex regions of primate genomes will be critical to realizing the full potential of these species as models of human genetic disease^74–76^.

## Materials and Methods

We sequenced and assembled eight NHP reference genomes using long-read PacBio HiFi and ONT sequencing chemistry and the hifiasm genome assembler^21^. All samples, with one exception, were female and correspond to the same samples used in previous studies as references, namely; Central chimpanzee (Clint)^7^, bonobo (Mhudiblu)^17^, Western gorilla (Kamilah)^13^, Sumatran orangutan (Susie)^8^, Northern white-cheeked gibbon (Asia)^10^, rhesus macaque (AG07107)^16^, common marmoset (CJ1700), and owl monkey (86718) (Table 1). We used pbsv, Sniffles, and PAV to characterize SVs and merged SVs using the SVPOP pipeline^37,38^. The merged calls were validated with HiFi sequencing data and assembly of select regions; ONT sequence data from the same specimens were used to calculate the false positive rate and validate assembly of select regions in our data set. The validated SVs were genotyped by Paragraph using Illumina WGS data from 120 population samples^16,39–43,84^. VEP was used to annotate the functional disruption of SVs^46^. In addition to SVs (<20 kbp) identified by the three callers, we used a graph-based aligner (Mashmap) to identify large structural changes across apes and Old World monkey^55^, defined here as SDRs. SDR validation was based on haplotype-resolved assemblies and ONT data. The ONT data also were used to call methylation by Guppy^85^. We also generated full-length Iso-Seq data specifically from 10 diverse marmoset tissues and from a gibbon immortalized lymphoblast line. In the case of the marmoset, full-length RNA was prepared from 10 distinct tissues obtained upon necropsy from a different specimen (*Callthrix jacchus*). Genomic divergence analyses were based on HiFi sequencing data and genomes, respectively. Syntenic regions across New World monkey to apes and MSAs were constructed with minimap2 and mafft^53,86^. The phylogenetic analyses were performed using TREEasy, IQTREE, and BEAST2^87–89^.

## Acknowledgments

We thank T. Brown for manuscript proofreading and editing. This article is subject to HHMI’s Open Access to Publications policy. HHMI lab heads have previously granted a nonexclusive CC BY 4.0 license to the public and a sublicensable license to HHMI in their research articles. Pursuant to those licenses, the author-accepted manuscript of this article can be made freely available under a CC BY 4.0 license immediately upon publication.

## Author contributions

Y.M. and E.E.E. conceived the project; Y.M., W.T.H., K.M.M., K.H., A.P.L., P.A.A., A.R., D.S.G., G.A.L., P.C.D., and E.E.E. generated sequencing data, assembled genomes, analyzed the data, and performed quality control analyses; X.Y., R.R., V.L.B., W.T.F., G.K.W., G.F., S.L.S., and W.C.W. contributed the marmoset and owl monkey samples; L.C. contributed the bonobo and gibbon samples; Y.M. performed the SNV divergence and ILS analyses; Y.M., W.T.H., P.A.A., S.Z., G.A.L., H.J., and E.E.E. performed SV analyses; Y.M. performed SDR analyses; M.H., and B.P. generated gene model annotations; Y.M., D.P., and E.E.E. performed *NPHP1* haplotype analyses; Y.M., X.W., and Q.L. performed the protein structure prediction analyses. Y.M. and E.E.E. drafted the manuscript. All authors read and approved the manuscript.

## Data and materials availability

The raw PacBio CLR, HiFi, and ONT data are deposited in NCBI under BioProject accession number PRJNA877605. The CLR and HiFi NHP assemblies are deposited in NCBI under BioProject accession numbers PRJNA941350-PRJNA941365. The marmoset Iso-Seq data are deposited under NCBI BioProject accession number PRJNA566173.

## References

1 Watson, J. D. The human genome project: past, present, and future. Science 248, 44–49, (1990).

2 Lander, E. S. et al. Initial sequencing and analysis of the human genome. Nature 409, 860–921, (2001).

3 Venter, J. C. et al. The sequence of the human genome. Science 291, 1304–1351 (2001).

4 Gibbs, R. A. The Human Genome Project changed everything. Nature reviews. Genetics 21, 575–576, doi:10.1038/s41576-020-0275-3 (2020).

5 Nurk, S. et al. The complete sequence of a human genome. Science 376, 44–53 (2022).

6 McConkey, E. H. et al. A primate genome project deserves high priority. Science 289, 1295–1296 (2000).

7 Chimpanzee Sequencing Analysis Consortium. Initial sequence of the chimpanzee genome and comparison with the human genome. Nature 437, 69–87 (2005).

8 Locke, D. P. et al. Comparative and demographic analysis of orang-utan genomes. Nature 469, 529–533 (2011).

9 Prüfer, K. et al. The bonobo genome compared with the chimpanzee and human genomes. Nature 486, 527–531 (2012).

10 Carbone, L. et al. Gibbon genome and the fast karyotype evolution of small apes. Nature 513, 195–201, (2014).

11 Rogers, J. & Gibbs, R. A. Comparative primate genomics: emerging patterns of genome content and dynamics. Nature Reviews Genetics 15, 347–359 (2014).

12 Juan, D., Santpere, G., Kelley, J. L., Cornejo, O. E. & Marques-Bonet, T. Current advances in primate genomics: novel approaches for understanding evolution and disease. Nature Reviews Genetics, 1–18 (2023).

13 Scally, A. et al. Insights into hominid evolution from the gorilla genome sequence. Nature 483, 169–175 (2012).

14 Gordon, D. et al. Long-read sequence assembly of the gorilla genome. Science 352, aae0344 (2016).

15 He, Y. et al. Long-read assembly of the Chinese rhesus macaque genome and identification of ape-specific structural variants. Nature communications 10, 4233 (2019).

16 Warren, W. C. et al. Sequence diversity analyses of an improved rhesus macaque genome enhance its biomedical utility. Science 370, eabc6617 (2020).

17 Mao, Y. et al. A high-quality bonobo genome refines the analysis of hominid evolution. Nature 594, 77–81 (2021).

18 Yang, C. et al. Evolutionary and biomedical insights from a marmoset diploid genome assembly. Nature 594, 227–233 (2021).

19 Kronenberg, Z. N. et al. High-resolution comparative analysis of great ape genomes. Science 360, eaar6343 (2018).

20 Logsdon, G. A., Vollger, M. R. & Eichler, E. E. Long-read human genome sequencing and its applications. Nature Reviews Genetics 21, 597–614 (2020).

21 Cheng, H., Concepcion, G. T., Feng, X., Zhang, H. & Li, H. Haplotype-resolved de novo assembly using phased assembly graphs with hifiasm. Nature methods 18, 170–175 (2021).

22 Rautiainen, M. et al. Telomere-to-telomere assembly of diploid chromosomes with Verkko. Nature Biotechnology, 1–9 (2023).

23 Mao, Y. & Zhang, G. A complete, telomere-to-telomere human genome sequence presents new opportunities for evolutionary genomics. Nature methods 19, 635–638 (2022).

24 Dennis, M. Y. et al. Evolution of human-specific neural SRGAP2 genes by incomplete segmental duplication. Cell 149, 912–922 (2012).

25 Fiddes, I. T. et al. Human-specific NOTCH2NL genes affect notch signaling and cortical neurogenesis. Cell 173, 1356–1369. e1322 (2018).

26 Kawanishi, K. et al. Human species-specific loss of CMP-N-acetylneuraminic acid hydroxylase enhances atherosclerosis via intrinsic and extrinsic mechanisms. Proceedings of the National Academy of Sciences 116, 16036–16045 (2019).

27 Logsdon, G. A. et al. The structure, function and evolution of a complete human chromosome 8. Nature 593, 101–107 (2021).

28 Mailund, T., Munch, K. & Schierup, M. H. Lineage sorting in apes. Annual review of genetics 48, 519–535 (2014).

29 Minh, B. Q., Hahn, M. W. & Lanfear, R. New methods to calculate concordance factors for phylogenomic datasets. Molecular biology and evolution 37, 2727–2733 (2020).

30 Steiper, M. E. & Young, N. M. Primate molecular divergence dates. Mol Phylogenet Evol 41, 384–394, (2006).

31 Wilkinson, R. D. et al. Dating primate divergences through an integrated analysis of palaeontological and molecular data. Systematic biology 60, 16–31, (2011).

32 Prado-Martinez, J. et al. Great ape genetic diversity and population history. Nature 499, 471–475, (2013).

33 Pozzi, L. et al. Primate phylogenetic relationships and divergence dates inferred from complete mitochondrial genomes. Mol Phylogenet Evol 75, 165–183, (2014).

34 de Manuel, M. et al. Chimpanzee genomic diversity reveals ancient admixture with bonobos. Science 354, 477–481, (2016).

35 Vanderpool, D. et al. Primate phylogenomics uncovers multiple rapid radiations and ancient interspecific introgression. PLoS biology 18, e3000954, (2020).

36 Álvarez-Carretero, S. et al. A species-level timeline of mammal evolution integrating phylogenomic data. Nature 602, 263–267, (2022).

37 Sedlazeck, F. J. et al. Accurate detection of complex structural variations using single-molecule sequencing. Nature methods 15, 461–468 (2018).

38 Ebert, P. et al. Haplotype-resolved diverse human genomes and integrated analysis of structural variation. Science 372, eabf7117 (2021).

39 Prado-Martinez, J. et al. Great ape genetic diversity and population history. Nature 499, 471–475 (2013).

40 Thomas, G. W. et al. Reproductive longevity predicts mutation rates in primates. Current Biology 28, 3193–3197. e3195 (2018).

41 Rogers, J. et al. The comparative genomics and complex population history of Papio baboons. Science Advances 5, eaau6947 (2019).

42 Okhovat, M. et al. Co-option of the lineage-specific LAVA retrotransposon in the gibbon genome. Proceedings of the National Academy of Sciences 117, 19328–19338 (2020).

43 Byrska-Bishop, M. et al. High-coverage whole-genome sequencing of the expanded 1000 Genomes Project cohort including 602 trios. Cell 185, 3426–3440. e3419 (2022).

44 Marques-Bonet, T. et al. A burst of segmental duplications in the genome of the African great ape ancestor. Nature 457, 877–881 (2009).

45 Sudmant, P. H. et al. Evolution and diversity of copy number variation in the great ape lineage. Genome research 23, 1373–1382 (2013).

46 McLaren, W. et al. The ensembl variant effect predictor. Genome biology 17, 1–14 (2016).

47 Moore, J. E. et al. Expanded encyclopaedias of DNA elements in the human and mouse genomes. Nature 583, 699–710 (2020).

48 Behesti, H. et al. ASTN2 modulates synaptic strength by trafficking and degradation of surface proteins. Proceedings of the National Academy of Sciences 115, E9717–E9726 (2018).

49 Bauleo, A. et al. Rare copy number variants in ASTN2 gene in patients with neurodevelopmental disorders. Psychiatric Genetics 31, 239–245 (2021).

50 Sentchordi-Montané, L. et al. Heterozygous aggrecan variants are associated with short stature and brachydactyly: description of 16 probands and a review of the literature. Clinical endocrinology 88, 820–829 (2018).

51 Efimova, E. V. et al. Trace amine-associated receptor 2 is expressed in the limbic brain areas and is involved in dopamine regulation and adult neurogenesis. Frontiers in Behavioral Neuroscience 16 (2022).

52 Chaisson, M. J. & Tesler, G. Mapping single molecule sequencing reads using basic local alignment with successive refinement (BLASR): application and theory. BMC bioinformatics 13, 1–18 (2012).

53 Li, H. Minimap2: pairwise alignment for nucleotide sequences. Bioinformatics 34, 3094–3100 (2018).

54 Porubsky, D. et al. Gaps and complex structurally variant loci in phased genome assemblies. bioRxiv, (2022).

55 Jain, C., Koren, S., Dilthey, A., Phillippy, A. M. & Aluru, S. A fast adaptive algorithm for computing whole-genome homology maps. Bioinformatics 34, i748–i756 (2018).

56 Yang, X. et al. A refined characterization of large-scale genomic differences in the first complete human genome. bioRxiv, (2022).

57 Sekar, A. et al. Schizophrenia risk from complex variation of complement component 4. Nature 530, 177–183 (2016).

58 Cantsilieris, S. et al. Recurrent structural variation, clustered sites of selection, and disease risk for the complement factor H (CFH) gene family. Proceedings of the National Academy of Sciences 115, E4433–E4442 (2018).

59 Thamadilok, S. et al. Human and nonhuman primate lineage-specific footprints in the salivary proteome. Molecular biology and evolution 37, 395–405 (2020).

60 Vollger, M. R. et al. Segmental duplications and their variation in a complete human genome. Science 376, eabj6965 (2022).

61 Jiang, Z. et al. Ancestral reconstruction of segmental duplications reveals punctuated cores of human genome evolution. Nature genetics 39, 1361–1368 (2007).

62 Khan, N. et al. Crystal structure of human PACRG in complex with MEIG1 reveals roles in axoneme formation and tubulin binding. Structure 29, 572–586. e576 (2021).

63 Zhang, Z. et al. MEIG1 is essential for spermiogenesis in mice. Proceedings of the National Academy of Sciences 106, 17055–17060 (2009).

64 Du, R. et al. Efficient typing of copy number variations in a segmental duplication-mediated rearrangement hotspot using multiplex competitive amplification. Journal of human genetics 57, 545–551 (2012).

65 Ciccarelli, F. D. et al. Complex genomic rearrangements lead to novel primate gene function. Genome research 15, 343–351 (2005).

66 Jumper, J. et al. Highly accurate protein structure prediction with AlphaFold. Nature 596, 583–589 (2021).

67 Parisi, M. A. et al. The NPHP1 gene deletion associated with juvenile nephronophthisis is present in a subset of individuals with Joubert syndrome. American journal of human genetics 75, 82–91, (2004).

68 Gana, S., Serpieri, V. & Valente, E. M. Genotype-phenotype correlates in Joubert syndrome: A review. Am J Med Genet C Semin Med Genet 190, 72–88, (2022).

69 Porubsky, D. et al. Recurrent inversion polymorphisms in humans associate with genetic instability and genomic disorders. Cell 185, 1986–2005. e1926 (2022).

70 Liao, W.-W. et al. A draft human pangenome reference. bioRxiv, 2022.2007. 2009.499321 (2022).

71 Wang, T. et al. The Human Pangenome Project: a global resource to map genomic diversity. Nature 604, 437–446 (2022).

72 Schneider, H. The current status of the New World monkey phylogeny. Anais da Academia Brasileira de Ciências 72, 165–172 (2000).

73 Perelman, P. et al. A molecular phylogeny of living primates. PLoS genetics 7, e1001342 (2011).

74 Baer, J. F., Weller, R. E. & Kakoma, I. Aotus: the owl monkey. (Academic Press, 2012).

75 Okano, H., Hikishima, K., Iriki, A. & Sasaki, E. in Seminars in fetal and neonatal medicine. 336–340 (Seminars in fetal and neonatal medicine).

76 Grillner, S. et al. Worldwide initiatives to advance brain research. Nature neuroscience 19, 1118–1122 (2016).

77 Nuttle, X. et al. Emergence of a Homo sapiens-specific gene family and chromosome 16p11. 2 CNV susceptibility. Nature 536, 205–209 (2016).

78 Hsieh, P. et al. Evidence for opposing selective forces operating on human-specific duplicated TCAF genes in neanderthals and humans. Nature communications 12, 5118 (2021).

79 Hsieh, P. et al. Adaptive archaic introgression of copy number variants and the discovery of previously unknown human genes. Science 366, eaax2083 (2019).

80 Ju, X.-C. et al. The hominoid-specific gene TBC1D3 promotes generation of basal neural progenitors and induces cortical folding in mice. Elife 5, e18197 (2016).

81 Dennis, M. Y. et al. The evolution and population diversity of human-specific segmental duplications. Nature ecology & evolution 1, 0069 (2017).

82 Prosser, S. L., Sahota, N. K., Pelletier, L., Morrison, C. G. & Fry, A. M. Nek5 promotes centrosome integrity in interphase and loss of centrosome cohesion in mitosis. Journal of Cell Biology 209, 339–348 (2015).

83 McAlear, T. S. & Bechstedt, S. The mitotic spindle protein CKAP2 potently increases formation and stability of microtubules. Elife 11, e72202 (2022).

84 Chen, S. et al. Paragraph: a graph-based structural variant genotyper for short-read sequence data. Genome biology 20, 1–13 (2019).

85 Liu, Y. et al. DNA methylation-calling tools for Oxford Nanopore sequencing: a survey and human epigenome-wide evaluation. Genome biology 22, 1–33 (2021).

86 Katoh, K. & Standley, D. M. MAFFT multiple sequence alignment software version 7: improvements in performance and usability. Molecular biology and evolution 30, 772–780 (2013).

87 Bouckaert, R. et al. BEAST 2: a software platform for Bayesian evolutionary analysis. PLoS computational biology 10, e1003537 (2014).

88 Nguyen, L.-T., Schmidt, H. A., Von Haeseler, A. & Minh, B. Q. IQ-TREE: a fast and effective stochastic algorithm for estimating maximum-likelihood phylogenies. Molecular biology and evolution 32, 268–274 (2015).

89 Mao, Y., Hou, S., Shi, J. & Economo, E. P. TREEasy: An automated workflow to infer gene trees, species trees, and phylogenetic networks from multilocus data. Mol Ecol Resour 20, (2020).

